# Mycorrhizal symbiosis alleviates plant water deficit within and across plant generations via plasticity

**DOI:** 10.1101/2020.07.21.213421

**Authors:** Javier Puy, Carlos P. Carmona, Inga Hiiesalu, Maarja Öpik, Francesco de Bello, Mari Moora

## Abstract

- Phenotypic plasticity is essential for organisms to adapt to local ecological conditions. It is expected that mutualistic interactions, such as arbuscular mycorrhizal (AM) symbiosis, mediate plant phenotypic plasticity, although it is not clear to what extent this plasticity may be heritable (i.e., transgenerational plasticity).
- We tested for plant plasticity within- and across-generations in response to AM symbiosis and varying water availability in a full factorial experiment over two generations, using a genetically uniform line of a perennial apomictic herb, *Taraxacum brevicorniculatum*. We examined changes in phenotype, performance, and AM fungal colonization of the offspring throughout plant development.
- AM symbiosis and water availability triggered phenotypic changes during the life cycle of plants. Additionally, both triggered adaptive transgenerational effects, especially detectable during the juvenile stage of the offspring. Water deficit and absence of AM fungi caused concordant plant phenotypic modifications towards a “stress-coping phenotype”, both within- and across-generations. AM fungal colonization of offspring was also affected by the parental environment.
- AM symbiosis can trigger within-generation and transgenerational plasticity in terms of functional traits related to resource-use acquisition and AM fungal colonization. Thus, transgenerational effects of mycorrhizal symbiosis are not limited to plant fitness, but also improve plants’ ability to cope with environmental stress.

## Introduction

Abiotic environment and prevailing biotic interactions select for the best-adapted individuals within and across species in terms of their functional traits (de Bello et al., 2012; Vellend, 2016). The probability of an individual to withstand this selection may depend on the nature and severity of the environmental condition, the heritable genetic variability, but also on its phenotypic plasticity (Price, Qvarnström, & Irwin, 2003). Phenotypic plasticity is the ability of an organism to modify its performance and trait expression in response to the environment without altering the DNA sequence (Price et al., 2003). These changes could operate within the life cycle of the organism subjected to those conditions (“within-generation plasticity”). Additionally, they may be transmitted to the following generations via transgenerational effects (also called “across-generation or transgenerational plasticity”), meaning that the abiotic and biotic environment experienced by the parental generation influence the phenotype of the offspring (Herman & Sultan, 2011; Jablonka & Raz, 2009). Recent studies suggest that transgenerational effects could play a key role in the adaptive response of organisms to stressors, proven particularly essential during the juvenile stages (Dechaine, Brock, & Weinig, 2015; Lämke & Bäurle, 2017; Latzel, Janeček, Doležal, Klimešová, & Bossdorf, 2014; Puy, Carmona, Dvořáková, Latzel, & de Bello, 2020). While the effect of abiotic conditions on transgenerational plasticity has been repeatedly demonstrated, little is known about the relative effect of transgenerational effects triggered by biotic conditions (Alonso, Ramos-Cruz, & Becker, 2019; Puy, de Bello, et al., 2020), and even less about how they interact with abiotic factors.

Together with species’ adaptations to environmental conditions in a site, biotic interactions are considered key drivers of plant community assembly (de Bello et al., 2012). Among these, positive interactions such as mycorrhizal symbiosis are essential in determining, and potentially expanding, the realized niches of species (Gerz, Bueno, Ozinga, Zobel, & Moora, 2018; Peay, 2016; van der Heijden, Wiemken, & Sanders, 2003). Arbuscular mycorrhizal (AM) symbiosis is a widespread mutualistic association between plant roots and fungi (from the subphylum Glomeromycotina; Spatafora et al., 2016). This association is considered mutually beneficial, since, in exchange for photosynthetic carbon, the AM fungi provide host plants with soil nutrients (mainly phosphates; Smith & Read, 2008), mitigate abiotic stress (e.g. making the host more tolerant to drought; Aroca, Porcel, & Ruiz-Lozano, 2012; Augé, Toler, & Saxton, 2015; Doubková, Vlasáková, & Sudová, 2013), and increase resistance to biotic stress, including pathogens (Pozo, López-Ráez, Azcón-Aguilar, & García-Garrido, 2015; Smith & Read, 2008). Besides the mutual effects of the AM interaction, the occurrence, abundance, activity, and the final outcome of the interaction (from positive to negative) are known to be affected by multiple factors (Hoeksema et al., 2010; N. C. Johnson, Graham, & Smith, 1997). These factors include intrinsic drivers such as the genotype of both partners, or the plant developmental stage or sex (Jones & Smith, 2004; Varga & Kytöviita, 2010). Also, external drivers such as the biotic environment (Šmilauer, Košnar, Kotilínek, & Šmilauerová, 2020) and/or soil nutrient and water availability (Martínez-García, de Dios Miranda, & Pugnaire, 2012; Pozo et al., 2015) might be important. Phosphorus, nitrogen or water deficiency in plants generally, at least when light is not limiting, stimulates AM symbiosis and influences the abundance of AM structures (i.e. arbuscules, vesicles, etc.; Martínez-García et al., 2012; Pozo et al., 2015). However, it remains unclear whether the environmental conditions experienced in parental generations could affect the functioning of AM symbiosis of the offspring generation (De Long et al., 2019).

As illustrated before, the fitness benefits of plants in AM symbiosis are relatively well known (Lu & Koide, 1994; Smith & Read, 2008). However, it is unclear whether these benefits partly operate through phenotypic plasticity induced by the interaction (Vannier, Mony, Bittebière, & Vandenkoornhuyse, 2015). AM symbiosis may lead to adaptive changes of plant morphological traits such as modifications in root architecture (Fusconi, 2014; Goh, Veliz Vallejos, Nicotra, & Mathesius, 2013; Nuortila, Kytöviita, & Tuomi, 2004) or in traits of the so called “plant economic spectrum”. Since these traits are associated with a fundamental trade-off between organisms along a resource-use-acquisition vs. resource-use-conservation gradient, they could be used to describe the plant resource-use strategy (Díaz et al., 2016). Moreover, it also remains unclear whether AM symbiosis in a parental generation can trigger similar or different phenotypic changes in the offspring (i.e. transgenerational effects) and whether these changes are beneficial (Koide, 2010; Varga, Vega-Frutis, & Kytöviita, 2013). As such, it is crucial to assess the relative effect of combined biotic and abiotic drivers on within-generation and transgenerational plasticity, as biotic drivers can potentially modulate the effect of abiotic stress (González, Dumalasová, Rosenthal, Skuhrovec, & Latzel, 2017; Metz, von Oppen, & Tielbörger, 2015).

It should be noted that transgenerational effects can be due to two mutually non-exclusive mechanisms: environmentally-induced heritable epigenetic modifications in the offspring (Lämke & Bäurle, 2017), or differences in seed provisioning, seed nutritional quality or hormonal balance provided by the maternal plants in the embryos (Dechaine et al., 2015; Germain, Grainger, Jones, & Gilbert, 2019; Herman & Sultan, 2011). In comparison, transgenerational effects originated from embryo modifications play more significant role during early stages of the development, but they tend to fade away with time (Latzel, Klimešová, Hájek, Gómez, & Šmilauer, 2010). Most of the existing evidence demonstrates that having mycorrhizal parents can be beneficial during the early stages of development of the offspring, i.e. increasing biomass, survival, growth rate, nutrient content, and seed production (Heppell, Shumway, & Koide, 1998; Koide, 2010; Varga, 2010; Varga et al., 2013). However, when testing for transgenerational effects of AM symbiosis, both mechanisms have rarely been considered (but see Varga et al. (2013) where seed mass was used as a covariate). Further, Varga & Soulsbury (2017, 2019) have investigated the potential epigenetic mechanism (i.e., DNA methylation) governing the transgenerational effects of AM symbiosis.**Error! Bookmark not defined.** Nevertheless, it has not yet been tested whether the transgenerational effects of AM symbiosis persist further into adult stage, neither whether these effects influence plant functional traits nor AM fungal colonization of the offspring.

We conducted a two-generation experiment to test for within- and across-generation plant plasticity in response to AM symbiosis using the perennial apomictic herb *Taraxacum brevicorniculatum*. Further, in order to test whether both plasticity types interact with abiotic stress conditions, we included a watering regime treatment. We then tested whether within- and across-generation plant plasticity were adaptive, e.g., resulting in an improved ability of the offspring to cope with water limitation. Additionally, we evaluated the persistence of the transgenerational effects throughout offspring development by measuring phenotypic traits, performance, and AM fungal colonization on juvenile and adult offspring.

## Materials and Methods

### Study material

*Taraxacum brevicorniculatum* Korol. is an obligate apomictic polycarpic perennial plant (Kirschner, Štěpánek, Černý, De Heer, & van Dijk, 2013). Like most species of the genus *Taraxacum*, it has a wide ecological niche, accepting all types of soils, pH and moisture levels (Luo & Cardina, 2012), and forms an active symbiosis with AM fungi (J. Puy, personal obs. based on a preliminary study). In this study we used genetically identical seeds collected from a population of plants grown under the same glasshouse conditions for several generations (collected and genetically identified by Kirschner *et al.* 2013). This strategy ensured homogeneous genetic and epigenetic variation in the plant material. Since *T. brevicorniculatum* is an obligate apomictic species, all seeds produced by a plant are effectively clones, thus enabling the study of plasticity within and across generations (Puy et al., 2018). In other words, all plants in the experiments were genetically identical, and after experiencing different conditions during the parental generation, the offspring only differed in non-genetic (i.e., non-genetic or epigenetic effects) inherited information.

### Experimental setup

#### Parental generation

In order to induce the potential transgenerational effects related to mycorrhizal symbiosis and water availability, we first conducted an experiment in which the parental generation was grown under different conditions in a glasshouse for three months (April-July 2017). We grew 364 genetically identical individuals of *T. brevicorniculatum* in individual pots (7 × 7 × 18 cm), half inoculated with AM fungi (AM) and the other half without (NM). In addition, half of these individuals were grown with sufficient water and half under a water deficit scenario, see below for details.

The substrate consisted of 2:1 mixture of sterilized sand and natural soil collected from a mesic meadow 30 km southeast of Tabor, 660 m a.s.l. (Vysočina region, Czech Republic, 49.331N, 15.003E) where *Taraxacum sect. Ruderalia* was present. The natural soil was firstly sieved using a 4 mm mesh sized sieve to remove any stones, macrofauna, or rhizomes from the soil. For the AM treatment the natural soil containing indigenous microbial community was used; whereas for the NM treatment the same soil was sterilized via γ irradiation (>25kGy dose; McNamara, Black, Beresford, & Parekh, 2003). To compensate for the loss of other soil microbes due to the sterilization, a microbial wash was also added (Liang et al., 2015). We obtained the microbial wash by blending 5kg of non-sterilized soil in 10 l water and filtering the solution through 20μm pore-size filter paper (Whatman® quantitative filter paper, Grade 41) broadly following van der Heijden *et al*., (1998). Gamma-sterilization did not change neither the C nor the N chemical composition of the soil compared to the non-sterilized soil (Fig. S1).

The AM and NM treatments were factorially combined with two levels of water availability. Half of the individuals were subjected to cycles of water deficit (simulating drought stress; W−), while the other half were watered regularly from the bottom ensuring the pot surface was always wet (control; W+). The water deficit treatment included periodic exclusion of watering until when 50% of the individuals had wilted leaves followed by one-week recovery in control conditions. By the end of the experiment, the water deficit treatment comprised two water deficit pulses (the first started 12^th^ of May and the second 15^th^ of June) that lasted three weeks plus one-week recovery each.

Prior to the establishment of the experiment, seeds were surface sterilised by immersion in 0.5% sodium hypochlorite solution (commercial bleach) for 20 minutes to avoid inoculation via seeds, and then germinated in Petri dishes. After 10 days of germination, the seedlings were transplanted individually into the pots specified above, with 91 replicates per treatment. After three months we harvested all the plants except 15 individuals from each of the four treatment combinations (AM W+, AM W−, NM W+, NM W−). These plants were kept for four more months in ambient conditions (water control condition), with a 12 h (20°C) / 12 h (10°C) light/darkness-and-temperature regime and with addition of fertilizer (Kristalon; NPK 15–5–30 + 3Mg + 5S) at the concentration of 300 ppm once per month, in order to promote seed production. Relocating the plants to more benign conditions was required to promote seed production especially for the non-mycorrhizal plants which, at the time of harvest, had not flowered yet. Then, seeds of each plant were collected, and after measuring the average seed mass per plant, were stored in cold (2-4 ºC).

#### Offspring experiment

A similar glasshouse experiment to the one described above was repeated the following year (April-August 2018) with the seeds produced by the parental generation in the first experiment. The aim of the offspring experiment was to test for adaptive transgenerational effects of AM symbiosis and water availability on the offspring at their juvenile and the adult stages. We tested this with a full factorial design where the offspring plants from each of the four parental treatments were exposed again to the four possible conditions (AM W+, AM W−, NM W+, NM W−). Thus, the offspring experimental design resulted in 16 combinations: two parental mycorrhizal inoculations (Par. M: AM/NM) x two parental water availability levels (Par. W: W+/W−) x two offspring mycorrhizal fungal inoculations (Off. M: AM/NM) x two offspring water availability levels (Off. W: W+/W−). Since the seed mass of AM parents was on average lower than that of NM parents (see below *Results of parental generation*, Table S1), and seed provisioning could be a potential mechanism of transgenerational effects (Herman & Sultan, 2011). We controlled for it by classifying seeds from all parental treatments into 5 size categories. Then, we took the same number of seeds from each size-group in each parental treatment, resulting in a similar distribution of seed sizes between parental treatments. Thus, the offspring experimental design finally resulted in the 16 combinations x 5 seed size categories x 8 seedlings = 640 pots (Fig. S2).

Plants were harvested at two different developmental stages. Half of the offspring plants were harvested 1.5 months after planting, at their juvenile stage; and the rest of the replicants were harvested five months after planting, at their adult stage before they flowered. Pots, substrate and watering regime were the same as in the parental experiment to ensure the most similar conditions. However, the first water deficit pulse in the offspring generation lasted four weeks instead of three (first one started the 25^th^ of April and the second one, the 1^st^ of June) to ensure comparable effects on plants physiology (i.e., percentage of plants with wilted leaves). To facilitate the application of the treatments, four replicates of a parental treatment were placed in parallel, one in each offspring treatment (Fig. S2).

### Measured traits

We measured a set of important plant traits in both generations. For each plant in the parental generation we measured survival, total dry biomass (aerial plus root biomass), seed output (i.e., number of seeds) and seed mass at the time of harvest. For five randomly chosen plants per treatment, we measured C, N and P content of the seeds, considered to be reliable indicators of seed reserve materials (Toorop et al., 2012). Total C and N content were determined by dry combustion using an elemental analyser (CHNS Elemental Analyzer vario MICRO cube, Elementar Analysensysteme GmbH, Germany). Total P was determined by flow injection analysis.

Additionally, we measured several above- and belowground vegetative plant traits. For each plant, two leaves were scanned for leaf area and weighed for fresh mass and dry mass after drying at 60º C (48h) to estimate specific leaf area (SLA; leaf area per dry mass, mm^2^/mg) and leaf dry matter content (LDMC; leaf dry mass per leaf fresh mass, mg/mg). Roots were carefully extracted, washed and a subsample of roots (6 cm^2^) was scanned at 600 dpi with an Epson Perfection 4990 scanner. Total root length, average root diameter (mm), and distribution of root length in different diameter classes were determined by using the image analysis software WinRHIZO Pro, 2008 (Regent Instruments Inc., Quebec, Canada). After scanning, the root subsample and the rest of the root system were dried for 48 h at 60 °C and weighed. We used these measurements to estimate specific root length (SRL; root length per dry mass, m/g), and fine roots percentage (root length with a diameter < 0.5mm per total root length). Further, we estimated root biomass allocation (i.e., root mass factor; RMF; root biomass per total biomass, g/g) after drying the remaining radicular part at 60º C (48h).

For each plant in the offspring generation, at the time of the respective harvest (i.e., juvenile and adult offspring harvest), we measured total dry biomass (aerial plus root biomass), and the same above- and belowground vegetative traits as described above. Additionally, we analyzed the content of C, N and P in the leaves of two randomly chosen plants from the juvenile stage and eight plants from the adult stage per treatment, following the methods described above. The root subsamples were stained with Chlorazol Black according to the protocol by Šmilauer, Košnar, Kotilínek, & Šmilauerová (2020). We quantified the AM fungal colonization by measuring the percentage of root length colonized (%RLC) by AM fungal structures (arbuscules, vesicles and hyphae). Magnified intersection method (McGonigle, Miller, Evans, Fairchild, & Swan, 1990) was used with 400 x magnification using a light microscope with graticule inserted into the eyepiece. All the specific structures of AM fungi (arbuscules, vesicles and hyphae) that intersected the vertical line (i.e., root in horizontal position) were counted for at least 100 intersections per root sample. We further calculated the arbuscule:vesicle ratio (relative abundance of arbuscules per vesicles), suggested as an indicator of the fungal activity status and the relative cost or benefit of the fungus to the host plant (Braunberger, Miller, & Peterson, 1991; Titus & Lepš, 2000).

### Statistical analysis

All analyses were carried out using R v3.2.3 (R Core Team 2016) with α=0.05 as significance level. In the parental generation, the effects of the mycorrhizal inoculation (two levels), the water availability (two levels), and their interaction were analysed by using linear effects models. Plant total biomass was always log-transformed.

In the offspring generation, individuals were grouped into sixteen different treatments (as a result of the combination of four factors with two levels each) depending on parental and offspring conditions. Two of the factors corresponded to parental conditions: mycorrhizal fungal inoculation treatment (Par. M: AM/NM), and water availability treatment (Par. W: W+/W−). The other two factors corresponded to offspring conditions: mycorrhizal fungal inoculation treatment (Off. M: AM/NM) and water availability treatment (Off. W: W+/W−). We analysed the effects of parental and offspring conditions on plant traits of the offspring using linear effects models where the four experimental factors (two parental, and two offspring conditions) and all their interactions were used as fixed effects. Additionally, in order to control for differences between parental treatments in seed provisioning or quality provided by the maternal plants (Herman & Sultan, 2011; Toorop et al., 2012), we included seed mass and seed stoichiometry as covariates in the model (i.e. as fixed effects). The seed stoichiometry values were computed assigning the scores of the first axis of a principal components analysis (PCA) that combined the C, N and P chemical composition of the seeds and absorbed 60% of the variation. Seed stoichiometry was not correlated with seed mass (Pearson correlation = −0.18, P = 0.44). Any effect of the parental conditions: either direct (Par. M or Par. W) or in interaction with the offspring conditions (Off. x Par.) that remained significant after removing the linear part of the maternal investment (seed mass and stoichiometry) was considered a transgenerational effect.

For the analysis of the effect of parental and offspring treatments on AM fungal colonization and arbuscule:vesicle ratio of the offspring, we used identical models, but excluding the offspring mycorrhizal inoculation factor (Off. M) from the model due to the lack of AM fungal colonization in the NM plants.

Additionally, we checked for correlations between plant traits and measures of AM fungal colonization to check which plant traits show plasticity in response to AM fungal colonization and to examine whether these changes could partially explain the benefits of AM symbiosis to the host plant, meaning that they are adaptive.

## Results

### Parental generation

In the parental generation, the water deficit treatment decreased *T. brevicorniculatum* total plant biomass and survival, but only on NM plants, with no effect on AM plants (Fig. S3a,b; Table S1). AM fungal inoculation increased plant growth, survival and reproductive investment (i.e., number of seeds per unit plant biomass), with no effect on the total number of seeds produced per plant (Fig. S3, Table S1). Additionally, seeds of AM plants were lighter than the ones of NM plants, and with higher N, P and C contents, although only the latter one was significant (Fig. S3, Table S1). Nevertheless, the seed macronutrients stoichiometry, i.e. C:N:P ratios, did not differ between AM and NM plants (Fig. S3, Table S1).

### Offspring plant traits

In the juvenile offspring, the majority of the measured plant traits were strongly affected by both offspring treatments (offspring mycorrhizal fungal inoculation treatment, Off. M; and offspring water availability treatment, Off. W), except leaf C:N ratio and SRL that were affected only by the mycorrhizal and by the water treatment respectively (Fig. 1 and Table S2). In general, plants under water deficit and absence of AM symbiosis were smaller, with higher biomass allocation into the roots (i.e., higher RMF), ticker leaves and roots (lower SLA and SRL), and lower P content and C:N ratio on leaves (Fig. 1 and Table S2).

**Figure 1:**
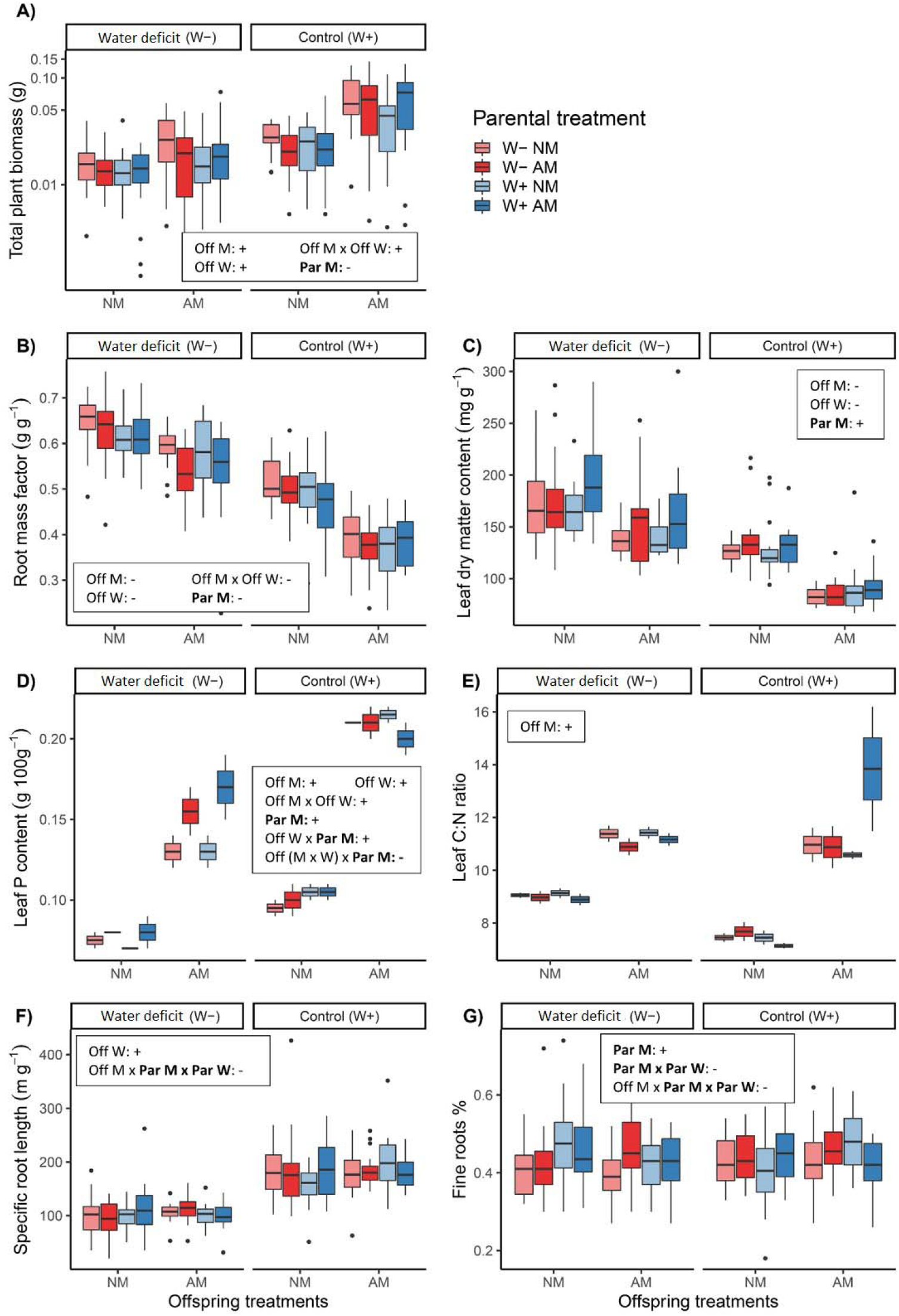
Effect of the offspring and parental treatments on plant phenotype characteristics of the juvenile offspring. a) total plant biomass, b) root mass factor, c) leaf dry matter content, d) leaf P content, e) leaf C:N ratio, f) specific root length and g) fine roots percentage. The significant factors of each model with the directionality of each effect are shown in the boxes. The factors corresponded to the offspring conditions: mycorrhizal inoculation treatment, Off. M; and water availability treatment Off. W; and the parental conditions (also highlighted in bold face): mycorrhizal inoculation treatment, Par. M; and water availability treatment, Par. W. Colour coding indicates the parental treatments: red - offspring of water-deficited parents, blue - offspring of parents that experienced water control conditions; intense colour - offspring of mycorrhizal parents, light colour - offspring of non-mycorrhizal parents. The bottom and top of the boxes are the 25th and 75th percentiles respectively, the centred band is the median and the whiskers represent 1.5 times the length of the box further from the box limits or the maximum or minimum observation in the absence of outliers.

After removing the effect of seed mass and seed stoichiometry (maternal investment effects), we found transgenerational effects (i.e., offspring plants were affected by the parental conditions) in most of the traits (i.e., seven out of nine, Table S2). When comparing the effects of both parental treatments (Par. M and Par. W), we found that parental mycorrhizal inoculation triggered transgenerational effects in all measured plant traits, while the parental water treatment induced changes only on three traits and generally in interaction with the offspring conditions (Fig. 1; Table S2). Juvenile offspring of mycorrhizal parents (Par. M) had in general lower biomass, lower allocation to the roots (lower RMF) but thinner roots (% Fine roots), and higher LDMC and leaf P content (Fig. 1a,b,c,d,g and Table S2). Except for biomass and LDMC, the direction of the response to the parental treatment was concordant with the response to the offspring treatment. For example, mycorrhizal offspring showed lower RMF and more P content on leaves, and these effects were further pronounced when offspring also had mycorrhizal parents (Fig. 1b,d and Table S2). Although in interaction with other factors, we found transgenerational effects induced by the parental water treatment. Mycorrhizal offspring of parents under water deficit (Par. W) increase the SRL and slim down the roots, when also had mycorrhizal parents (Par. W x Off. M x Par. M Fig. 1f,g and Table S2).

At the adult stage, the offspring treatments (Off. M and Off. W) were still the main drivers of plant plasticity (Fig. 2 and Table S3) and generally the responses of the traits were in the same direction than those during juvenile stage (leaf P content and total biomass; Fig. 2 and Table S3). Nonetheless, LDMC plasticity in response to offspring conditions (Off. M and Off. W) reversed compared with the juvenile stage. The offspring with water deficit and absence of AM symbiosis had higher LDMC during the juvenile stage, but lower LDMC during the adult stage (Fig. 1b and Fig. 2b). Similar reversed response happened in RMF only when offspring were with water deficit. Also, root traits that did not respond to the offspring treatments during juvenile stage, started responding during adult stage. At the latter stage, plants with water deficit and non-mycorrhizal plants had thinner and more absorptive roots (higher SRL and % Fine roots; Fig. 2f, g and Table S3). Finally, at this stage, we still detected significant transgenerational effects on plant traits, but only on RMF (Fig. 2b and Table S3). Adult offspring of mycorrhizal parents (Par. M) still had lower allocation to the roots (lower RMF).

**Figure 2:**
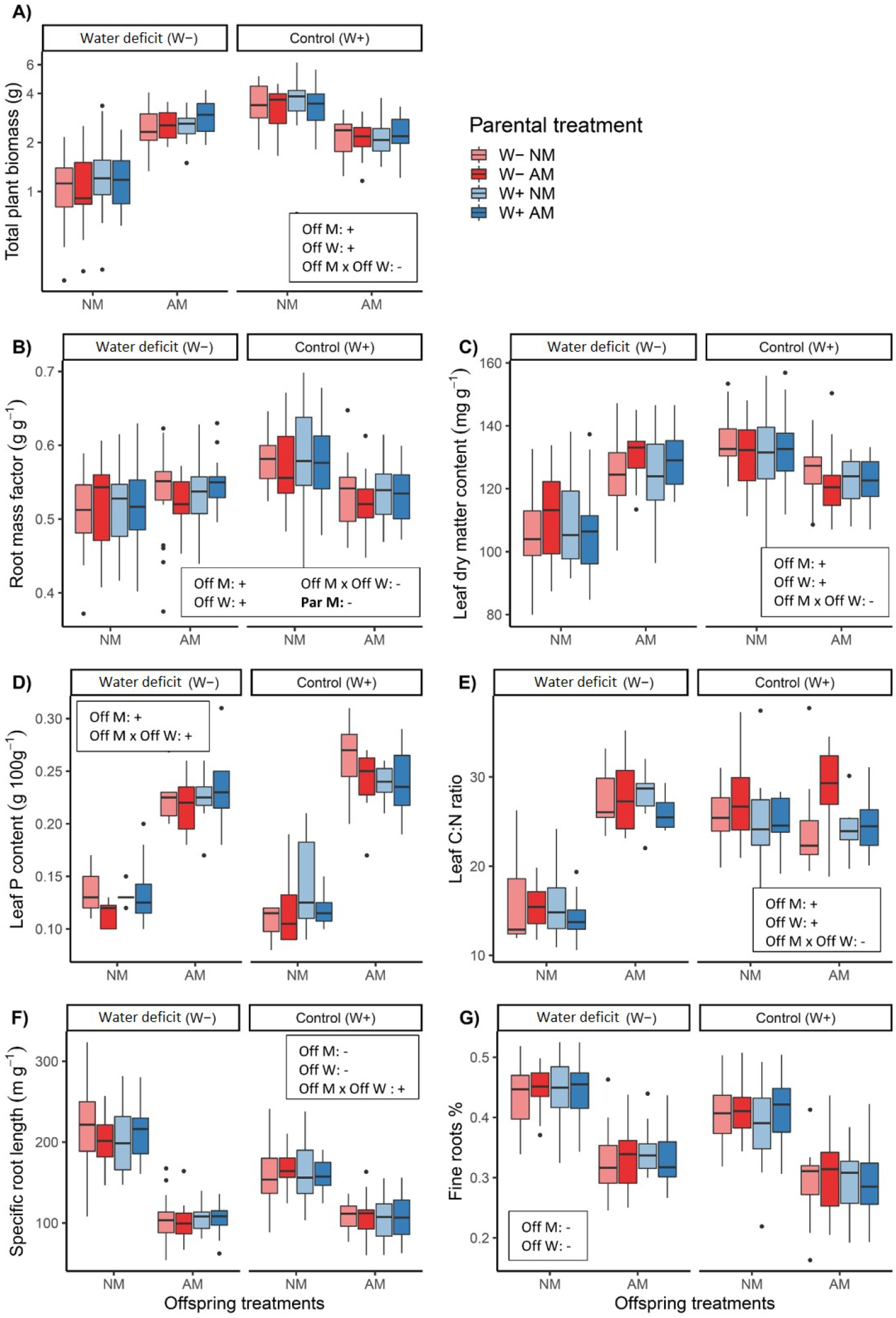
Effect of the offspring and parental treatments on plant phenotype characteristics of the adult offspring. a) total plant biomass, b) root mass factor, c) leaf dry matter content, d) leaf P content, e) leaf C:N ratio, f) specific root length and g) fine roots percentage. The significant factors of each model with the directionality of each effect are shown in the boxes. The factors corresponded to the offspring conditions: mycorrhizal fungal inoculation treatment, Off. M; and water availability treatment Off. W; and the parental conditions (also highlighted in bold face): mycorrhizal inoculation treatment, Par. M; and water availability treatment, Par. W. Colour coding indicates the parental treatments: red - offspring of water-deficited parents, blue - offspring of parents that experienced water control conditions; intense colour - offspring of mycorrhizal parents, light colour - offspring of non-mycorrhizal parents.

### Offspring AM fungal colonization

The water availability treatments (Off. W) modified the AM fungal colonization of the juvenile offspring. Plants grown with water deficit cycles had less root length colonized by AM fungi (lower %RLC) but had higher arbuscule:vesicle ratio than control plants (Fig. 3a, b and Table S2). Additionally, AM fungal colonization was affected by the parental treatments only when offspring plants had water deficit (Off. W x Par. W; Table S2). Offspring of parents that did not experience water deficit had higher %RLC than the ones of parents that had experienced water deficit (Fig. 3a; Table S2). For the arbuscule:vesicle ratio we did not detect significant transgenerational effects (Fig. 3b, Table S2 and Fig. S4).

**Figure 3:**
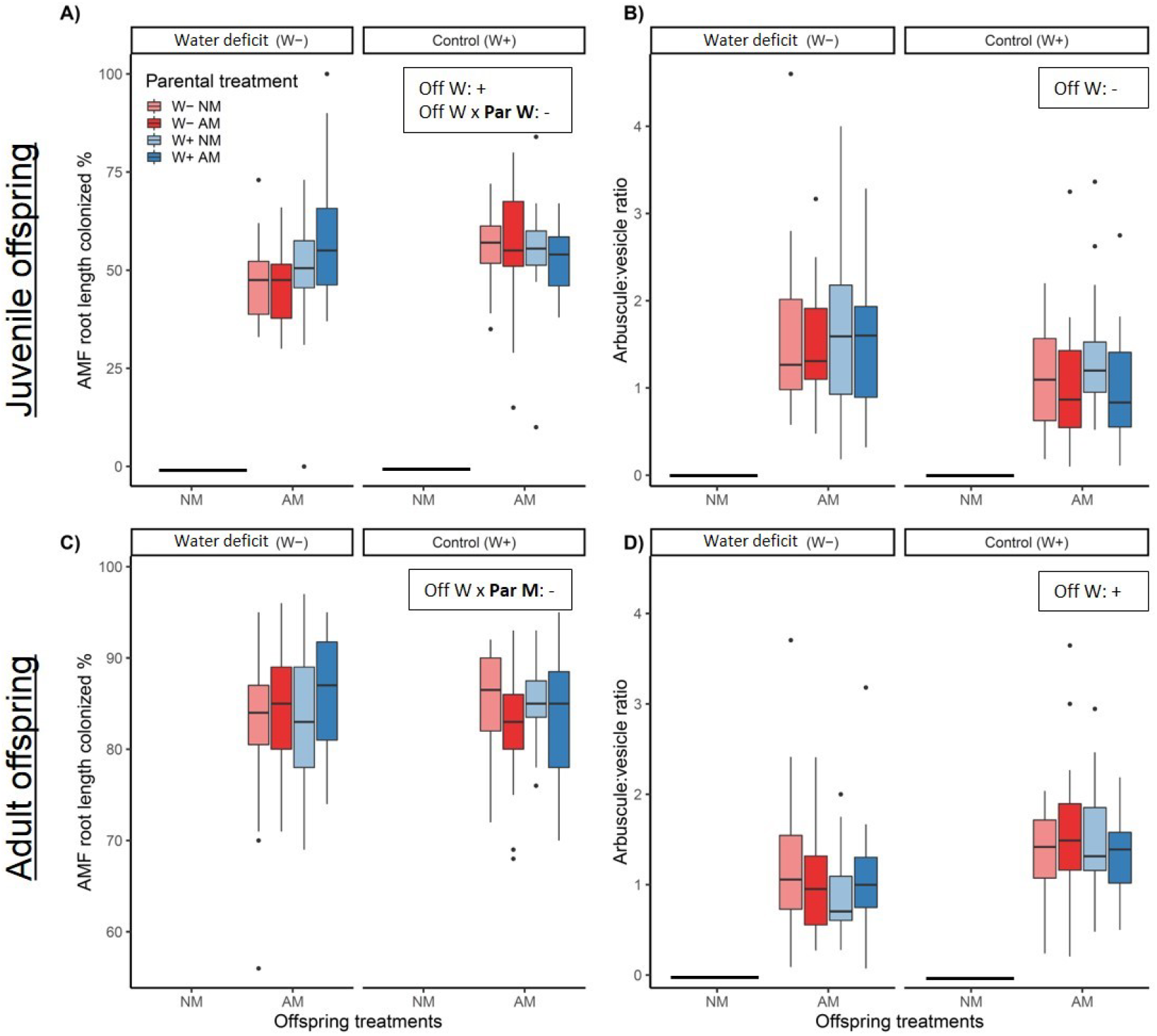
Effect of the offspring and parental treatments on AM fungal root colonisation in juvenile stage (upper row) and adult stage (lower row): a) and c) percentage of root length colonized by AM fungi; b) and d) arbuscule:vesicle ratio. The significant factors of each model with the directionality of each effect are shown in the boxes. The factors corresponded to the offspring conditions: mycorrhizal inoculation treatment, Off. M; and water availability treatment Off. W; and the parental conditions (also highlighted in bold face): mycorrhizal inoculation treatment, Par. M; and water availability treatment, Par. W. Colour coding indicates the parental treatments: red - offspring of water-deficited parents, blue - offspring of parents that experienced water control conditions; intense colour - offspring of mycorrhizal parents, light colour - offspring of non-mycorrhizal parents. The nonmycorrhizal offspring treatment plants were not colonized by AM fungi.

During the adult stage, we found no significant difference in %RLC between the offspring water availability treatments (Off. W), although there was a lower arbuscule:vesicle ratio in plants with water deficit, reversed response compared with what happened during the juvenile stage (Fig. 3c, d, Table S3 and Fig. S4). Also, at this stage we found that AM fungal colonization was affected by the parental treatments only when the offspring plants grew with water deficit cycles. The adult offspring of mycorrhizal parents (Off. W x Par. M) had higher %RLC than the ones of non-mycorrhizal parents (Fig 3c; Table S3).

## Discussion

In this study we show the importance of AM symbiosis in triggering phenotypic plasticity in plants, both during their life cycle and in following generations. Such phenotypic changes can improve the ability of individuals to cope with environmental stress and likely increase the species’ realized niche. Here we show that mycorrhizal symbiosis could specifically trigger morphological changes related to resource use and resource acquisition strategies within-generations and to succeeding generations. Further, we provide evidence that AM fungal colonization of the offspring could be also affected by parental conditions. Transgenerational effects of mycorrhizal symbiosis and water availability were not caused by differences in the quality and resources provided in the seed (Herman & Sultan, 2011), pointing to heritable epigenetic mechanisms as potential factor transmitting and mediating these effects across generations.

### Within-generational plasticity on offspring traits is development specific

The strong response of plants of *T. brevicorniculatum* to the conditions experienced during their life cycle (i.e., offspring mycorrhizal fungal inoculation and water availability treatments) shows the high level of plasticity of this plant species. However, we found that the response to these conditions differed in juvenile and adult phases, suggesting specific plant plasticity at different developmental stages (Coleman, McConnaughay, & Ackerly, 1994).

As expected, measurements of different fitness-related characteristics (i.e. plant nutrition and growth) suggest that AM symbiosis improved plant performance and mitigated water deficit, since the benefit of being mycorrhizal increased with decreasing water supply (Aroca et al., 2012; Augé et al., 2015; Doubková et al., 2013). First, mycorrhizal fungal inoculation dramatically increased leaf P content at both developmental stages of the offspring (Fig. 1d and Fig. 2d). These results reflect that mycorrhizal plants had better nutritional supply – were provided with important nutrients such as P (Doubková et al., 2013; Lu & Koide, 1994) – despite being in water deficit conditions. Probably because of this, mycorrhizal fungal inoculation also increased growth (i.e., plant biomass) of plant individuals in water deficit conditions to similar levels than growth of non-stress plants. However, this benefit was more pronounced in adult offspring than in juveniles (Fig. 2a vs Fig. 1a), probably due to a greater cost/benefit ratio during the juvenile stage (N. C. Johnson et al., 1997).

In both developmental stages, mycorrhizal fungal inoculation and water availability treatments induced significant changes in multiple plant phenotypic traits related to the resource-use strategy of the plant (also called plant economic spectrum; Díaz et al., 2016), including both below- and aboveground traits (Fusconi, 2014; Goh et al., 2013; Nuortila et al., 2004). However, traits seem to respond differently depending on the developmental stage of the plant. Plant phenotypic plasticity triggered by AM fungal inoculation could be either direct (i.e., caused in reaction to AM fungal infection), or indirect by a lack of phenotypic plasticity in reaction to water deficit, since the AM fungi increase the tolerance of host plants to it (Goh et al., 2013; Maherali, 2014).

During the juvenile stage water availability and mycorrhizal fungal inoculation triggered independent and additive trait plasticity in the same direction (Table S2). Similar to findings of Shumway & Koide (1994), under reduced water availability and/or absence of AM fungi, plants shifted towards more resource-conservative phenotype based on the plant economics spectrum framework (Díaz et al., 2016). This phenotype is characterized by having greater belowground biomass allocation (RMF), less photosynthetic but more water-use efficient leaves (greater LDMC and C:N ratio, lower SLA) and thicker and more resistant roots (lower SRL) (Fig. 1 and Fig. S5). These traits are expected to be beneficial when resources are scarce, since they are associated with longer lifespan and enhance water use efficiency of the plant under water stress (Díaz et al., 2016). Thus, the plastic response towards a conservative phenotype could improve *T. brevicorniculatum* ability to cope with water deficit. In this case, AM symbiosis seems to induce direct plant trait plasticity, and not indirectly by a lack of reaction to water deficit. Accordingly, the host plant with more root length colonized by AM fungal (%RLC), had thinner roots, and higher SLA (Fig. S5a).

During the adult stage, the direction of plasticity changed compared to the response at the juvenile stage. In response to reduced water availability and/or absence of AM fungi plants shifted towards more resource-acquisitive phenotype. The plants decreased LDMC and C:N ratio and increased their SRL and percentage of fine roots, reflecting an adaptive phenotypic plasticity that improved resource uptake and compensated for the lack of AM symbiosis and the involvement of extraradical mycelium in plant resource uptake (Fusconi, 2014; Goh et al., 2013; Pozo et al., 2015). At this stage, the plasticity triggered by AM symbiosis seems to be indirect and caused by a lack of physiological reaction of the plant to water deficit (i.e., it induced the same phenotypic changes as the water control condition), because AM symbiosis have increased plant tolerance against stress. This different responses depending on developmental stage could explain why previous studies found variable effects of mycorrhizal symbiosis on plant phenotype (D. Johnson, Martin, Cairney, & Anderson, 2012; Nuortila et al., 2004). Nevertheless, at both stages we found strong positive correlation between root length colonized by AM fungal (%RLC) and SRL and strong negative one between %RLC and root diameter (Fig. S5b).

### Transgenerational effects on offspring performance and phenotype

Several studies had already shown that offspring from mycorrhizal parents can have greater fitness, reflected in higher biomass, survival, growth rate, and seed production (Heppell et al., 1998; Koide, 2010; Varga et al., 2013). We partially confirmed this in this study. We found that offspring of mycorrhizal parents had higher leaf P content than those of non-mycorrhizal parents (Fig. 1d), suggesting that mycorrhizal symbiosis, besides directly providing soil nutrients to host plants, could also increase offspring’s nutrient uptake via transgenerational effects. However, in terms of biomass, we observed that indeed having non-mycorrhizal parents was beneficial (Fig. 1a). This contrasting result in biomass was probably due to the fact that, unlike previous evidence, we experimentally after controlling the effect of seed mass and stoichiometry (Dechaine et al., 2015; Germain et al., 2019; Herman & Sultan, 2011; but see Varga & Kytöviita, 2010). Also it could have further allowed us to detected underlying transgenerational effects probably controlled by epigenetic or hormonal mechanisms (Herman & Sultan, 2011; Rottstock, Kummer, Fischer, & Joshi, 2017; Varga & Soulsbury, 2017). One way to confirm whether these effects were only epigenetically controlled would be to modify the epigenetic signature of the plants via application of a demethylation agent (Puy, de Bello, et al., 2020; Puy et al., 2018). However, it should be first tested whether the demethylation application does not harm the AM fungal community.

Moreover, we found transgenerational effects in offspring traits linked with the resource use and exploitation strategy of the plant (Díaz et al., 2016), with transgenerational plasticity of the offspring being concordant with the within-generational response to the treatments. Plant traits shifted towards a “stress-coping phenotype under water deficit and absence of AM symbiosis during their life cycle (i.e., more conservative phenotype: increased the RMF and decrease SRL). Additionally, offspring individuals became even more conservative when their parents were under those stressful conditions (i.e., water deficit and absence of AM symbiosis; Fig. 1). Thus, the transgenerational effects reinforced trait plasticity when individuals where grown in the same conditions of their parents. This suggests a “stress memory” effect that could improve the ability of plants to cope with predictable environment across generations. The higher biomass found in offspring of non-mycorrhizal parents could partly be a consequence of this adaptive heritable phenotypic plasticity, that enhance their water use efficiency despite the environment they grow on.

Even though both abiotic and biotic parental environments seemed to trigger transgenerational effects, we found that mycorrhizal fungal inoculation affected plant traits more than water availability (Fig. 1 and Table S2). Since *T. brevicorniculatum* has a wide ecological niche – it grows well under different water availability conditions (Luo & Cardina, 2012) – it is likely that water is a less crucial stressor for this species, and likewise that the transgenerational effects due to water availability could have been not evolutionary nor ecological strongly constraining (Rendina González, Preite, Verhoeven, & Latzel, 2018). Also, it is important to emphasize that we found that transgenerational effects on phenotype were expressed early on the ontogeny (Fig. 1) and faded away over offspring life development (Fig. 2). This result reinforces the idea of transgenerational effects as an important factor promoting adaptation to repeated ecological conditions, especially during juvenile stages and establishment of communities (Dechaine et al., 2015; Latzel et al., 2014). Also, even though transgenerational effects are likely to fade away with time, the effects associated to differences in the seeds might fade away even faster (Latzel et al., 2010) as happened in our case (Table S2 and S3).

### Within- and transgenerational effect on AM fungal colonization

As expected, the environmental factors experienced by offspring during their life cycle (Off W) affected the offspring AM fungal colonization. Contrary to our expectation (Martínez-García et al., 2012; Pozo et al., 2015), water deficiency did not stimulate root AM fungal colonization (since %RLC decreased). However, the proportion of arbuscules:vesicles increased in offspring with water deficit, suggesting that water deficiency can stimulate AM fungal activity, since the arbuscules are the main structures where resource exchange takes place.

Moreover, we found that AM fungal colonization of the offspring could be influenced by the conditions experienced by the host plant parental generation. To our knowledge, this is the first study that shows transgenerational effects on AM fungal colonization (but see De Long et al., 2019 for differences in AMF structures). Specifically, we observed that only under water deficit conditions, the offspring of parents that experienced water scarcity had lower AM fungal colonization. Although this appears to contradict our initial hypothesis that resource scarcity (i.e., water deficit and NM parental treatments) would stimulate AM symbiosis of the offspring, we must note that offspring from water deficit and NM parents compensated by having increased root biomass. This means that this offspring had in total more roots colonized and greater number of arbuscules per individual than offspring from parents under ambient water conditions (see the calculations made by relativizing to the total root biomass of the plant – i.e., root biomass x %RLC – Fig. S6). Thus, we conclude that the result in general still supports the notion that transgenerational effects modify offspring towards the “stress-coping phenotype” stimulating the establishment and activity of the AM symbiosis.

As also found for plant traits, the AM fungal colonization of the offspring was influenced by the parental conditions still during adult stage. This suggests that transgenerational effects could influence plant–AM fungi relationship and persist further than just the establishment and early stages of the symbiosis. At the adult stage, %RLC was affected by the parental mycorrhizal status, so that offspring from mycorrhizal parents had higher %RLC under water deficit. These results suggest that mycorrhizal symbiosis could be promoted in the offspring when the parental generation had experienced mycorrhizal symbiosis.

## Conclusions

We found that mycorrhizal symbiosis, alone and in combination with water availability, triggered phenotypic changes within and across generations on plant performance and AM fungal colonization. Water deficits and absence of AM fungi triggered concordant plant phenotypic plasticity, towards a stress-coping phenotype, both within- and across generations. This reflects an adaptive epigenetic mechanism that promotes rapid adaptation, and probably improves the ability of the species to cope with water deficit. In a context where the importance of individual and intraspecific variation of mycorrhizal plants and fungi in ecosystems is increasingly acknowledged (D. Johnson et al., 2012), our study provides evidence of a mechanism of variation that has been neglected so far: transgenerational plasticity. These plastic changes confer competitive advantages to the next generation. Importantly, we show that transgenerational effects remained significant even after controlling by the differences in seed provisioning and nutritional quality stocked by the mother, pointing to other mechanisms such as heritable epigenetic mechanisms as potential mediators and transmitters of these effects across generations.

## Acknowledgments

We thank M. Applová, T. Jairus and N. G. Medina for technical assistance and N. Plowman for English revision. This research was financially supported by the Czech Science Foundation grant GACR (Grant No. GA17-11281S) and the European Union through the European Regional Development Fund. IH, MÖ, CPC and MM received support by grants from the Estonian Research Council (PSG293, PUT1170) and by the European Regional Development Fund (Centre of Excellence EcolChange).

## Author contribution

JP, CPC, IH, MÖ, FdB and MM designed the research; JP performed the experiments; JP and CPC analysed the data; JP wrote the main manuscript. All authors contributed substantially to revisions and gave final approval for publication.

## Data availability

The data that support the findings of this study will be available on Figshare repository with acceptance.

## Supporting information

**Fig. S1**: N and C content of the substrate (AMF and NM).

**Fig. S2**: Schematic representation of the design of the offspring experiment.

**Fig. S3**: Effect of the treatments on parental generation.

**Fig. S4**: Effect of the offspring and parental treatments on frequency of AM structures of the juvenile and adult offspring.

**Fig. S5**: Correlation between pairs of plant traits and AM fungal colonization measured in juvenile and adult offspring plants.

**Fig. S6**: Relative effect of the offspring and parental treatments on total root AM fungal colonization and percentage of arbuscules on juvenile offspring.

**Table S1**: Summary of the linear mixed-effect model for main and interaction effects of the treatments in the parental generation.

**Table S2**: Summary of the linear mixed-effect model for main and interaction effects of offspring and parental treatments on juvenile offspring.

**Table S3**: Summary of the linear mixed-effect model for main and interaction effects of offspring and parental treatments on adult offspring.

## References

Alonso, C., Ramos-Cruz, D., & Becker, C. (2019). The role of plant epigenetics in biotic interactions. New Phytologist, 221(2), 731–737. doi: 10.1111/nph.15408

Aroca, R., Porcel, R., & Ruiz-Lozano, J. M. (2012). Regulation of root water uptake under abiotic stress conditions. Journal of Experimental Botany, 63(1), 43–57. doi: 10.1093/jxb/err266

Augé, R. M., Toler, H. D., & Saxton, A. M. (2015). Arbuscular mycorrhizal symbiosis alters stomatal conductance of host plants more under drought than under amply watered conditions: a meta-analysis. Mycorrhiza, 25(1), 13–24. doi: 10.1007/s00572-014-0585-4

Braunberger, P. G., Miller, M. H., & Peterson, R. L. (1991). Effect of phosphorus nutrition on morphological characteristics of vesicular=arbuscular mycorrhizal colonization of maize. New Phytologist, 119(1), 107–113. doi: 10.1111/j.1469-8137.1991.tb01013.x

Coleman, J. S., McConnaughay, K. D. M., & Ackerly, D. D. (1994). Interpreting phenotypic variation in plants. Trends in Ecology & Evolution, 9(5), 187–191. doi: 10.1016/0169-5347(94)90087-6

de Bello, F., Price, J. N., Münkemüller, T., Liira, J., Zobel, M., Thuiller, W.,…Pärtel, M. (2012). Functional species pool framework to test for biotic effects on community assembly. Ecology, 93(10), 2263–2273. doi: 10.1890/11-1394.1

De Long, J. R., Semchenko, M., Pritchard, W. J., Cordero, I., Fry, E. L., Jackson, B. G.,…Bardgett, R. D. (2019). Drought soil legacy overrides maternal effects on plant growth. Functional Ecology, 00, 1–11. doi: 10.1111/1365-2435.13341

Dechaine, J. M., Brock, M. T., & Weinig, C. (2015). Maternal environmental effects of competition influence evolutionary potential in rapeseed (Brassica rapa). Evolutionary Ecology, 29(1), 77–91. doi: 10.1007/s10682-014-9735-6

Díaz, S., Kattge, J., Cornelissen, J. H. C., Wright, I. J., Lavorel, S., Dray, S.,…Gorné, L. D. (2016). The global spectrum of plant form and function. Nature, 529(7585), 167–171. doi: 10.1038/nature16489

Doubková, P., Vlasáková, E., & Sudová, R. (2013). Arbuscular mycorrhizal symbiosis alleviates drought stress imposed on Knautia arvensis plants in serpentine soil. Plant and Soil, 370(1–2), 149–161. doi: 10.1007/s11104-013-1610-7

Fusconi, A. (2014). Regulation of root morphogenesis in arbuscular mycorrhizae: What role do fungal exudates, phosphate, sugars and hormones play in lateral root formation? Annals of Botany, 113(1), 19–33. doi: 10.1093/aob/mct258

Germain, R. M., Grainger, T. N., Jones, N. T., & Gilbert, B. (2019). Maternal provisioning is structured by species’ competitive neighborhoods. Oikos, 128(1), 45–53. doi: 10.1111/oik.05530

Gerz, M., Bueno, C. G., Ozinga, W. A., Zobel, M., & Moora, M. (2018). Niche differentiation and expansion of plant species are associated with mycorrhizal symbiosis. Journal of Ecology, 106(1), 254–264. doi: 10.1111/1365-2745.12873

Goh, C.-H., Veliz Vallejos, D. F., Nicotra, A. B., & Mathesius, U. (2013). The Impact of Beneficial Plant-Associated Microbes on Plant Phenotypic Plasticity. Journal of Chemical Ecology, 39(7), 826–839. doi: 10.1007/s10886-013-0326-8

González, A. P. R., Dumalasová, V., Rosenthal, J., Skuhrovec, J., & Latzel, V. (2017). The role of transgenerational effects in adaptation of clonal offspring of white clover (Trifolium repens) to drought and herbivory. Evolutionary Ecology, 31(3), 345–361. doi: 10.1007/s10682-016-9844-5

Heppell, K. B., Shumway, D. L., & Koide, R. T. (1998). The effect of mycorrhizal infection of Abutilon theophrasti on competitiveness of offspring. Functional Ecology, 12(2), 171–175. doi: 10.1046/j.1365-2435.1998.00188.x

Herman, J. J., & Sultan, S. E. (2011). Adaptive Transgenerational Plasticity in Plants: Case Studies, Mechanisms, and Implications for Natural Populations. Frontiers in Plant Science, 2, 102. Retrieved from http://journal.frontiersin.org/article/10.3389/fpls.2011.00102/abstract

Hoeksema, J. D., Chaudhary, V. B., Gehring, C. A., Johnson, N. C., Karst, J., Koide, R. T.,…Umbanhowar, J. (2010). A meta-analysis of context-dependency in plant response to inoculation with mycorrhizal fungi. Ecology Letters, 13(3), 394–407. doi: 10.1111/j.1461-0248.2009.01430.x

Jablonka, E., & Raz, G. (2009). Transgenerational Epigenetic Inheritance: Prevalence, Mechanisms, and Implications for the Study of Heredity and Evolution. The Quarterly Review of Biology, 84(2), 131–176. doi: 10.1086/598822

Johnson, D., Martin, F., Cairney, J. W. G., & Anderson, I. C. (2012). The importance of individuals: Intraspecific diversity of mycorrhizal plants and fungi in ecosystems. New Phytologist, 194(3), 614–628. doi: 10.1111/j.1469-8137.2012.04087.x

Johnson, N. C., Graham, J. H., & Smith, F. A. (1997). Functioning of mycorrhizal associations along the mutualism-parasitism continuum. New Phytologist, 135(4), 575–586. doi: 10.1046/j.1469-8137.1997.00729.x

Jones, M. D., & Smith, S. E. (2004). Exploring functional definitions of mycorrhizas: Are mycorrhizas always mutualisms? Canadian Journal of Botany, 82(8), 1089–1109. doi: 10.1139/b04-110

Kirschner, J., Štěpánek, J., Černý, T., De Heer, P., & van Dijk, P. J. (2013). Available ex situ germplasm of the potential rubber crop Taraxacum koksaghyz belongs to a poor rubber producer, T. brevicorniculatum (Compositae-Crepidinae). Genetic Resources and Crop Evolution, 60(2), 455–471. doi: 10.1007/s10722-012-9848-0

Koide, R. T. (2010). Mycorrhizal Symbiosis and Plant Reproduction. In Arbuscular Mycorrhizas: Physiology and Function (pp. 297–320). doi: 10.1007/978-90-481-9489-6_14

Lämke, J., & Bäurle, I. (2017, December 27). Epigenetic and chromatin-based mechanisms in environmental stress adaptation and stress memory in plants. Genome Biology, Vol. 18, p. 124. doi: 10.1186/s13059-017-1263-6

Latzel, V., Janeček, Š., Doležal, J., Klimešová, J., & Bossdorf, O. (2014). Adaptive transgenerational plasticity in the perennial Plantago lanceolata. Oikos, 123(1), 41–46. doi: 10.1111/j.1600-0706.2013.00537.x

Latzel, V., Klimešová, J., Hájek, T., Gómez, S., & Šmilauer, P. (2010). Maternal effects alter progeny’s response to disturbance and nutrients in two Plantago species. Oikos, 119(11), 1700–1710. doi: 10.1111/j.1600-0706.2010.18737.x

Liang, M., Liu, X., Etienne, R. S., Huang, F., Wang, Y., & Yu, S. (2015). Arbuscular mycorrhizal fungi counteract the Janzen-Connell effect of soil pathogens. Ecology, 96(2), 562–574. doi: 10.1890/14-0871.1

Lu, X., & Koide, R. T. (1994). The effects of mycorrhizal infection on components of plant growth and reproduction. New Phytologist, 128(2), 211–218. Retrieved from https://nph.onlinelibrary.wiley.com/doi/pdf/10.1111/j.1469-8137.1994.tb04004.x

Luo, J., & Cardina, J. (2012). Germination patterns and implications for invasiveness in three Taraxacum (Asteraceae) species. Weed Research, 52(2), 112–121. doi: 10.1111/j.1365-3180.2011.00898.x

Maherali, H. (2014). Is there an association between root architecture and mycorrhizal growth response? New Phytologist, 204(1), 192–200. doi: 10.1111/nph.12927

Martínez-García, L. B., de Dios Miranda, J., & Pugnaire, F. I. (2012). Impacts of changing rainfall patterns on mycorrhizal status of a shrub from arid environments. European Journal of Soil Biology, 50, 64–67. doi: 10.1016/J.EJSOBI.2011.12.005

McGonigle, T. P., Miller, M. H., Evans, D. G., Fairchild, G. L., & Swan, J. A. (1990). A new method which gives an objective measure of colonization of roots by vesicular—arbuscular mycorrhizal fungi. New Phytologist, 115(3), 495–501. doi: 10.1111/j.1469-8137.1990.tb00476.x

McNamara, N. P., Black, H. I. J., Beresford, N. A., & Parekh, N. R. (2003). Effects of acute gamma irradiation on chemical, physical and biological properties of soils. Applied Soil Ecology, 24(2), 117–132. doi: 10.1016/S0929-1393(03)00073-8

Metz, J., von Oppen, J., & Tielbörger, K. (2015). Parental environmental effects due to contrasting watering adapt competitive ability, but not drought tolerance, in offspring of a semi-arid annual Brassicaceae. Journal of Ecology, 103(4), 990–997. doi: 10.1111/1365-2745.12411

Nuortila, C., Kytöviita, M.-M., & Tuomi, J. (2004). Mycorrhizal symbiosis has contrasting effects on fitness components in Campanula rotundifolia. New Phytologist, 164(3), 543–553. doi: 10.1111/j.1469-8137.2004.01195.x

Peay, K. G. (2016). The Mutualistic Niche: Mycorrhizal Symbiosis and Community Dynamics. Annual Review of Ecology, Evolution, and Systematics, 47(1), 143–164. doi: 10.1146/annurev-ecolsys-121415-032100

Pozo, M. J., López-Ráez, J. A., Azcón-Aguilar, C., & García-Garrido, J. M. (2015, March 1). Phytohormones as integrators of environmental signals in the regulation of mycorrhizal symbioses. New Phytologist, Vol. 205, pp. 1431–1436. doi: 10.1111/nph.13252

Price, T. D., Qvarnström, A., & Irwin, D. E. (2003, July 22). The role of phenotypic plasticity in driving genetic evolution. Proceedings of the Royal Society B: Biological Sciences, Vol. 270, pp. 1433–1440. doi: 10.1098/rspb.2003.2372

Puy, J., Carmona, C. P., Dvořáková, H., Latzel, V., & de Bello, F. (2020). Diversity of parental environments increases phenotypic variation in Arabidopsis populations more than genetic diversity but similarly affects productivity. Annals of Botany, mcaa100. doi: 10.1093/aob/mcaa100

Puy, J., de Bello, F., Dvořáková, H., Medina, N. G., Latzel, V., & Carmona, C. P. (2020). Competition-induced transgenerational plasticity influences competitive interactions and leaf decomposition of offspring. New Phytologist, nph.17037. doi: 10.1111/nph.17037

Puy, J., Dvořáková, H., Carmona, C. P., de Bello, F., Hiiesalu, I., & Latzel, V. (2018). Improved demethylation in ecological epigenetic experiments: Testing a simple and harmless foliar demethylation application. Methods in Ecology and Evolution, 9(3), 744–753. doi: 10.1111/2041-210X.12903

Rendina González, A. P., Preite, V., Verhoeven, K. J. F., & Latzel, V. (2018). Transgenerational Effects and Epigenetic Memory in the Clonal Plant Trifolium repens. Frontiers in Plant Science, 9, 1677. doi: 10.3389/fpls.2018.01677

Rottstock, T., Kummer, V., Fischer, M., & Joshi, J. (2017). Rapid transgenerational effects in Knautia arvensis in response to plant community diversity. Journal of Ecology, 105(3), 714–725. doi: 10.1111/1365-2745.12689

Shumway, D. L., & Koide, R. T. (1994). Reproductive responses to mycorrhizal colonization of Abutilon theophrasti Medic, plants grown for two generations in the field. New Phytologist, Vol. 128, pp. 219–224. doi: 10.1111/j.1469-8137.1994.tb04005.x

Šmilauer, P., Košnar, J., Kotilínek, M., & Šmilauerová, M. (2020). Contrasting effects of host identity, plant community, and local species pool on the composition and colonization levels of arbuscular mycorrhizal fungal community in a temperate grassland. New Phytologist, 225(1), 461–473. doi: 10.1111/nph.16112

Smith, S. E., & Read, D. J. (2008). Mycorrhizal symbiosis. Academic Press.

Spatafora, J. W., Chang, Y., Benny, G. L., Lazarus, K., Smith, M. E., Berbee, M. L.,…Stajich, J. E. (2016). A phylum-level phylogenetic classification of zygomycete fungi based on genome-scale data. Mycologia, 108(5), 1028–1046. doi: 10.3852/16-042

Titus, J. H., & Lepš, J. (2000). The response of arbuscular mycorrhizae to fertilization, mowing, and removal of dominant species in a diverse oligotrophic wet meadow. American Journal of Botany, 87(3), 392–401. doi: 10.2307/2656635

Toorop, P. E., Campos Cuerva, R., Begg, G. S., Locardi, B., Squire, G. R., & Iannetta, P. P. M. (2012). Co-adaptation of seed dormancy and flowering time in the arable weed Capsella bursa-pastoris (shepherds purse). Annals of Botany, 109(2), 481–489. doi: 10.1093/aob/mcr301

van der Heijden, M. G. A., Klironomos, J. N., Ursic, M., Moutoglis, P., Streitwolf-Engel, R., Boller, T.,…Sanders, I. R. (1998). Mycorrhizal fungal diversity determines plant biodiversity, ecosystem variability and productivity. Nature, 396(6706), 69–72. doi: 10.1038/23932

van der Heijden, M. G. A., Wiemken, A., & Sanders, I. R. (2003). Different arbuscular mycorrhizal fungi alter coexistence and resource distribution between co-occurring plant. New Phytologist, 157(3), 569–578. doi: 10.1046/j.1469-8137.2003.00688.x

Vannier, N., Mony, C., Bittebière, A. K., & Vandenkoornhuyse, P. (2015). Epigenetic mechanisms and microbiota as a toolbox for plant phenotypic adjustment to environment. Frontiers in Plant Science, 6(DEC), 1159. doi: 10.3389/fpls.2015.01159

Varga, S. (2010). Effects of arbuscular mycorrhizas on reproductive traits in sexually dimorphic plants: a review. Spanish Journal of Agricultural Research, 8(S1), 11. doi: 10.5424/sjar/201008s1-5299

Varga, S., & Kytöviita, M. M. (2010). Mycorrhizal benefit differs among the sexes in a gynodioecious species. Ecology, 91(9), 2583–2593. doi: 10.1890/09-1383.1

Varga, S., & Soulsbury, C. D. (2017). Paternal arbuscular mycorrhizal fungal status affects DNA methylation in seeds. Biology Letters, 13(9), 20170407. doi: 10.1098/rsbl.2017.0407

Varga, S., & Soulsbury, C. D. (2019). Arbuscular mycorrhizal fungi change host plant DNA methylation systemically. Plant Biology, 21(2), 278–283. doi: 10.1111/plb.12917

Varga, S., Vega-Frutis, R., & Kytöviita, M. M. (2013). Transgenerational effects of plant sex and arbuscular mycorrhizal symbiosis. New Phytologist, 199(3), 812–821. doi: 10.1111/nph.12305

Vellend, M. (2016). The Theory of Ecological Communities (MPB-57). In The Theory of Ecological Communities (MPB-57). doi: 10.1515/9781400883790

