## Supplementary matterial for "Mycorrhizal symbiosis alleviates plant water deficit within and across plant generations via plasticity" (from another version of this preprint: version 1, 2020-07-22)

**Supporting Information:**

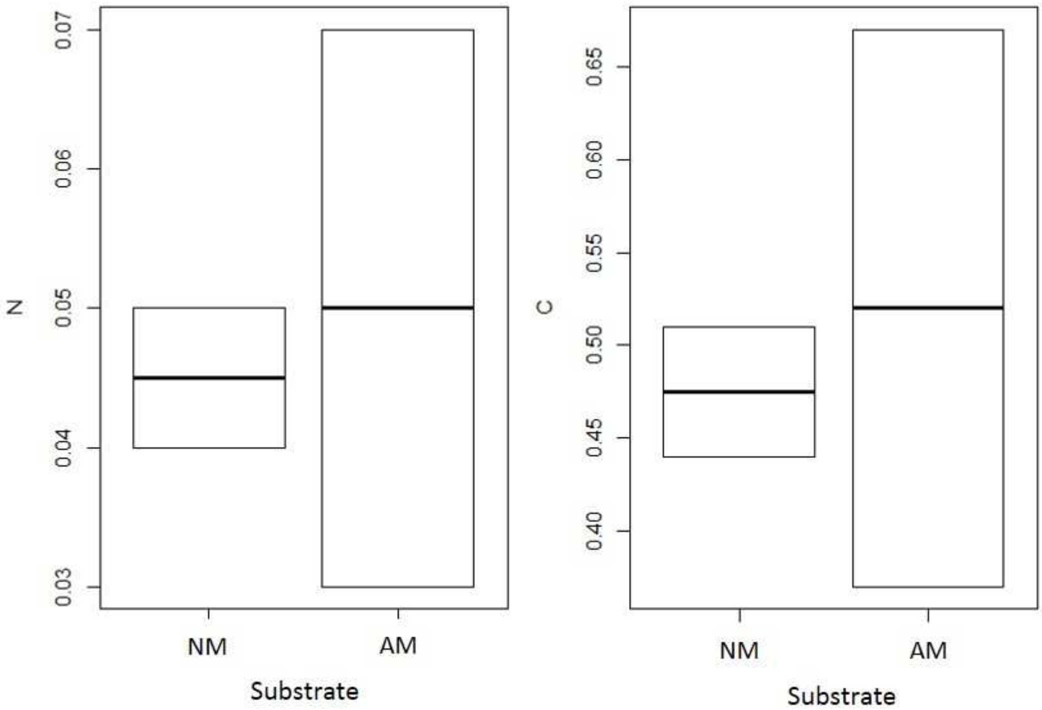


**Figure S1**: N and C content of the sterilized substrate (NM) and substrate with AMF (AM).

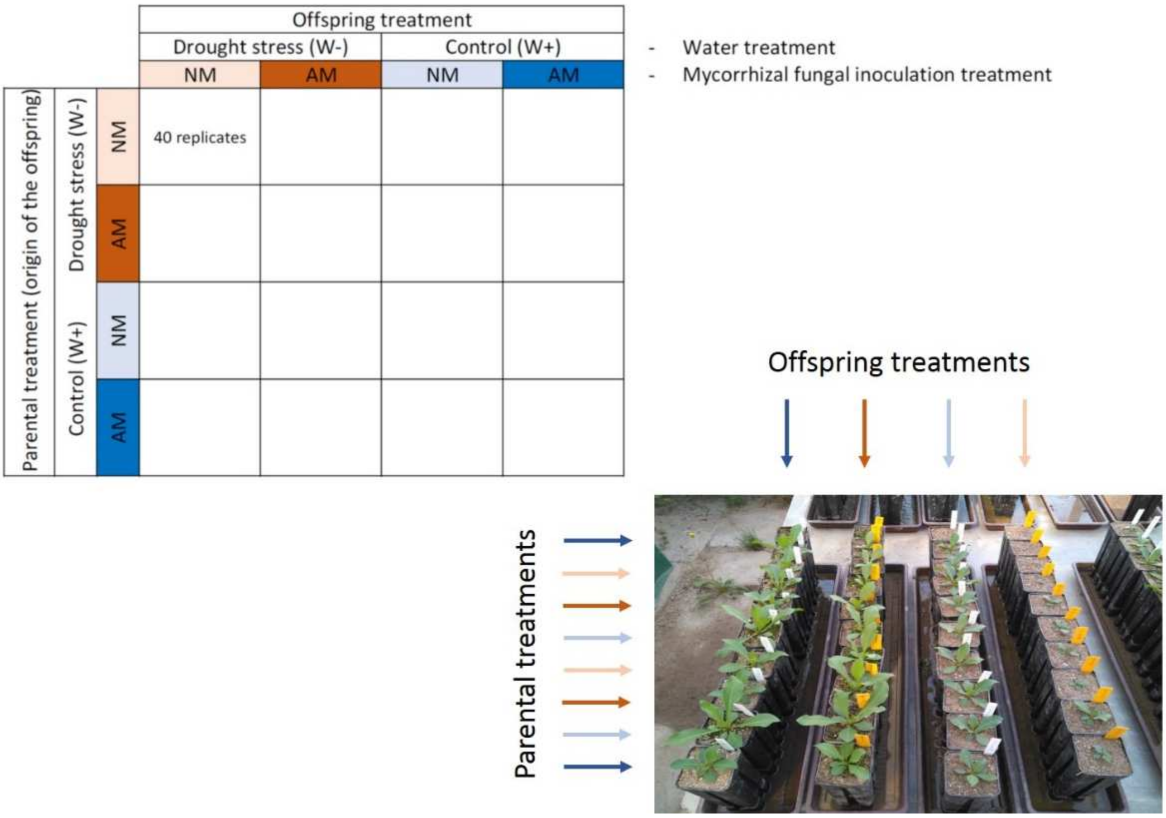


**Figure S2**: Schematic representation of the design of the offspring experiment, and detailed representation of the experimental blocks.

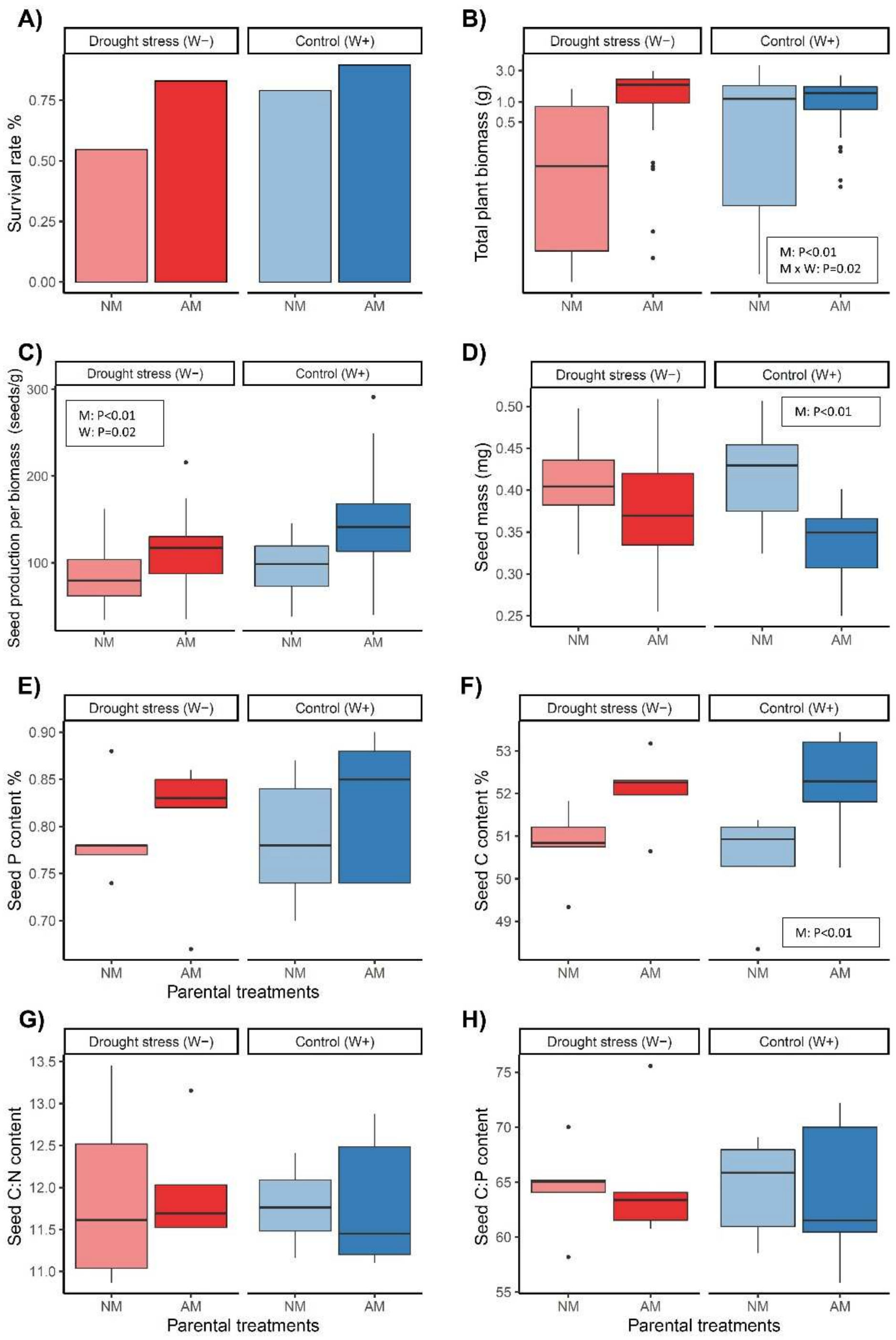


**Figure S3**: Effect of the treatments on parental generation: a) survival rate, b) total plant biomass, c) number of seeds produced per biomass, d) average seed mass, e) seed P content, f) seed C content, g) seed C:N ratio and h) seed C:P ratio. The significant factors of each model are shown in the boxes. The different coloured boxplots represent the treatments. The bottom and top of the boxes are the 25th and 75th percentiles respectively, the centred band is the median and the whiskers represent 1.5 times the length of the box further from the box limits or the maximum or minimum observation in the absence of outliers.

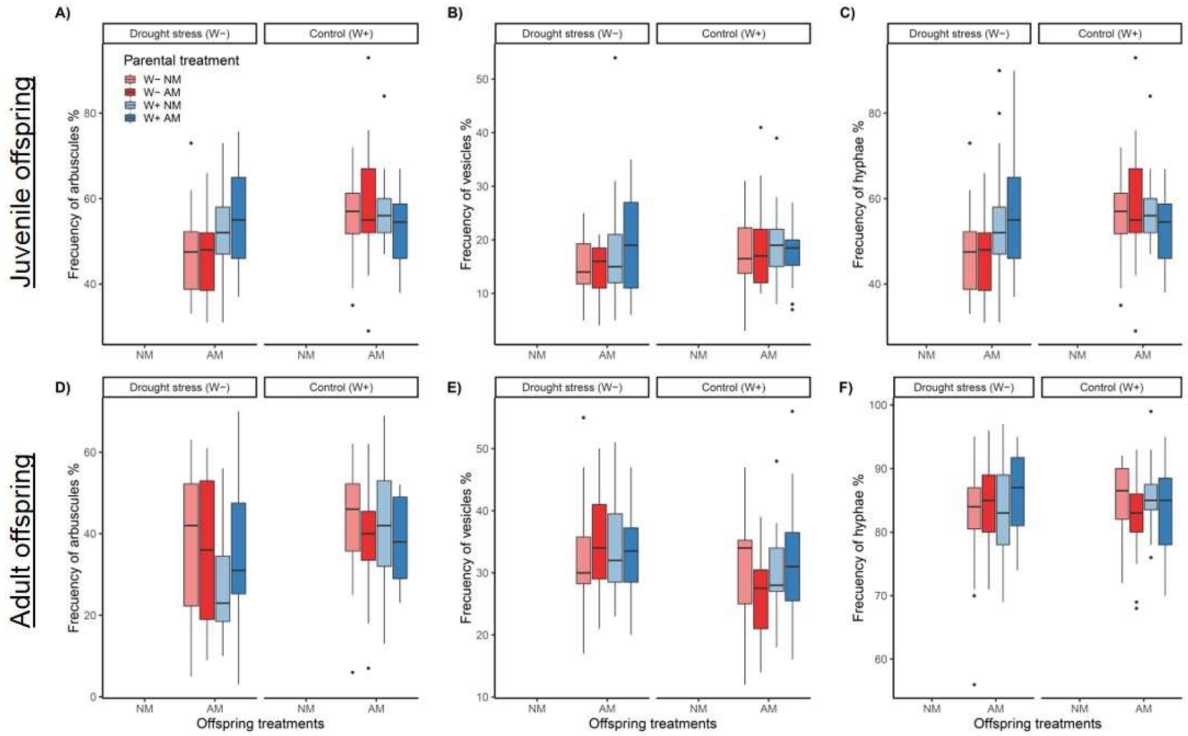


**Figure S4**: Effect of the offspring and parental treatments on other AM symbiosis characteristics of the juvenile offspring (upper row) and adult offspring (lower row): a) and d) frequency of arbuscules b) and e) vesicles and c) and f) hyphae. In each of the offspring treatments, the different coloured boxplots represent the parental treatments: offspring of water-stressed parents in red and offspring of parents that experienced water control conditions in blue; and offspring of mycorrhizal parents more intense coloured, and offspring of non-mycorrhizal parents with a lighter colour. The nonmycorrhizal offspring treatment plants were not AMF colonized. The bottom and top of the boxes are the 25th and 75th percentiles respectively, the centred band is the median and the whiskers represent 1.5 times the length of the box further from the box limits or the maximum or minimum observation in the absence of outliers.

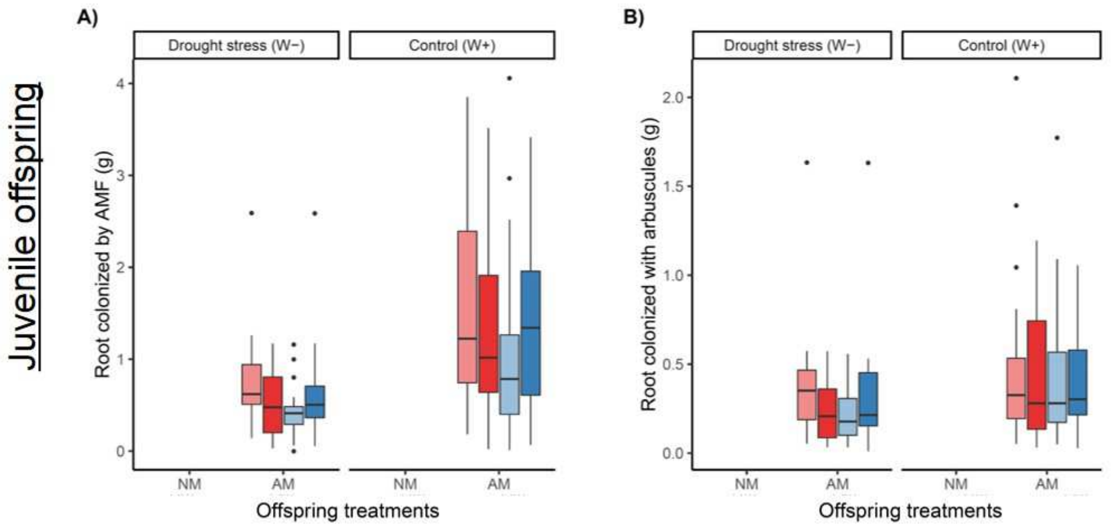


**Figure S5:** Effect of the offspring and parental treatments on total AM fungal root colonisation in juvenile stage relativized by the individual root biomass: a) total mass of root colonized by AM fungi; and b) total mass of root colonized with arbuscules. There are no significant effects of the parental factors.

### Tables:

**Table S1: Summary of the linear mixed-effect model for main and interaction effects of the treatments in the parental generation.** Degrees of freedom followed by F values and P values are given for all the effects analysed. Significant results are shown in bold face, and the colour indicates the direction of the effect (positive in blue, negative in red)

| Source of variation | df | log (Total biomass) | | Seed production | | Seed production per plant biomass | | Seed mass | | Seed stoichiometry | | | | Seed macronutrient content | | | | | |
|---|---|---|---|---|---|---|---|---|---|---|---|---|---|---|---|---|---|---|---|
| | | | | | | | | | | C:N ratio | | C:P ratio | | C content | | N content | | P content | |
| | | F | P | F | P | F | P | F | P | F | P | F | P | F | P | F | P | F | P |
| Mycorrhizas inoculation treatment | 1 | 36.8 | **<0.01** | 0.4 | 0.51 | 11.7 | **<0.01** | 20.5 | **<0.01** | 0.03 | 0.86 | 0.00 | 0.98 | **9.7** | **<0.01** | 0.73 | 0.41 | 0.68 | 0.42 |
| Water availability treatment | 1 | 3.4 | 0.06 | 0.0 | 0.99 | 5.3 | **0.02** | 1.9 | 0.17 | 0.15 | 0.70 | 0.05 | 0.83 | 0.06 | 0.81 | 0.06 | 0.80 | 0.04 | 0.85 |
| Mycorrhizas x Water | 1 | 5.2 | **0.02** | 0.0 | 0.92 | 0.7 | 0.38 | 1.9 | 0.17 | 0.00 | 0.95 | 0.04 | 0.84 | 0.25 | 0.62 | 0.11 | 0.74 | 0.10 | 0.75 |

**Table S2: Summary of the linear mixed-effect model for main and interaction effects of offspring and parental treatments (the latter highlighted in bold) on the juvenile offspring.** Degrees of freedom followed by χ2 values and P values are given for all the effects analysed. Significant results are shown in bold, and the colour indicates the direction of the effect (positive in blue, negative in red)

| | | Plant traits | | | | | | | | | | | | | | | | | | AMF colonization | | | |
|---|---|---|---|---|---|---|---|---|---|---|---|---|---|---|---|---|---|---|---|---|---|---|---|
| | | Whole-plant traits | | | | Leaf traits | | | | | | | | Root traits | | | | | | | | | |
| Source of variation | df | Total biomass | | Root mass factor (RMF) | | Leaf dry matter content (LDMC) | | Specific leaf area (SLA) | | Leaf P content | | Leaf C:N content | | Specific root length (SRL) | | Average root diameter | | % Fine roots | | Root length colonized by AMF (%RLC) | | Arbuscules: vesicles ratio | |
| | | χ2 | P | χ2 | P | χ2 | P | χ2 | P | χ2 | P | χ2 | P | χ2 | P | χ2 | P | χ2 | P | χ2 | P | χ2 | P |
| Seed mass | 1 | 2.7 | 0.102 | 3.2 | 0.073 | 0.0 | 0.973 | 3.1 | 0.078 | 0.2 | 0.663 | 0.0 | 0.865 | 2.9 | 0.088 | 1.4 | 0.238 | 0.2 | 0.636 | 0.6 | 0.448 | 2.4 | 0.123 |
| Offspring Mycorrhizal inoculation (Off. M) | 1 | 68.2 | **<0.01** | 123.6 | **<0.01** | 136.1 | **<0.01** | 335.3 | **<0.01** | 354.5 | **<0.01** | 82.9 | **<0.01** | 0.4 | 0.503 | 5.6 | **0.018** | 0.0 | 0.967 | | | | |
| Offspring Water availability (Off. W) | 1 | 137.7 | **<0.01** | 401.0 | **<0.01** | 320.3 | **<0.01** | 286.4 | **<0.01** | 86.1 | **<0.01** | 3.1 | 0.076 | 226.1 | **<0.01** | 57.8 | **<0.01** | 0.3 | 0.594 | 4.6 | **0.032** | 11.5 | **<0.01** |
| **Parental Mycorrhizal inoculation (Par. M)** | 1 | 1.1 | 0.301 | 7.5 | **<0.01** | 9.5 | **<0.01** | 1.3 | 0.246 | 2.7 | 0.102 | 0.2 | 0.690 | 0.1 | 0.708 | 0.5 | 0.498 | 1.3 | 0.260 | 0.3 | 0.594 | 1.5 | 0.221 |
| **Parental Water availability (Par. W)** | 1 | 1.3 | 0.262 | 0.5 | 0.464 | 0.6 | 0.444 | 2.4 | 0.121 | 0.1 | 0.776 | 0.7 | 0.403 | 0.3 | 0.615 | 0.4 | 0.552 | 0.4 | 0.522 | 1.6 | 0.209 | 1.5 | 0.225 |
| Off. M x Off. W | 1 | 21.7 | **<0.01** | 10.1 | **<0.01** | 5.9 | **0.015** | 58.8 | **<0.01** | 15.8 | **<0.01** | 7.7 | **<0.01** | 0.8 | 0.365 | 11.0 | **<0.01** | 2.2 | 0.141 | | | | |
| **Par. M x Par. W** | 1 | 1.1 | 0.301 | 0.4 | 0.521 | 0.5 | 0.480 | 0.1 | 0.760 | 0.1 | 0.759 | 0.4 | 0.537 | 0.0 | 0.854 | 0.9 | 0.338 | 3.2 | 0.076 | 0.1 | 0.785 | 1.4 | 0.243 |
| Off. M x **Par. M** | 1 | 2.9 | 0.091 | 0.2 | 0.687 | 0.0 | 0.990 | 0.7 | 0.403 | 0.6 | 0.426 | 1.0 | 0.309 | 0.5 | 0.497 | 0.0 | 0.965 | 0.5 | 0.475 | | | | |
| Off. M x **Par. W** | 1 | 0.2 | 0.618 | 0.7 | 0.419 | 0.1 | 0.759 | 1.6 | 0.208 | 0.0 | 1.000 | 1.5 | 0.217 | 0.1 | 0.749 | 0.0 | 0.879 | 0.6 | 0.427 | | | | |
| Off. W x **Par. M** | 1 | 0.1 | 0.700 | 1.9 | 0.170 | 4.6 | **0.032** | 0.1 | 0.808 | 5.7 | **0.017** | 2.3 | 0.133 | 0.4 | 0.514 | 0.4 | 0.535 | 0.0 | 0.961 | 0.5 | 0.478 | 0.1 | 0.793 |
| Off. W x **Par. W** | 1 | 0.2 | 0.656 | 0.0 | 0.852 | 0.2 | 0.623 | 1.2 | 0.277 | 0.0 | 1.000 | 0.4 | 0.534 | 0.4 | 0.547 | 4.1 | **0.043** | 2.7 | 0.102 | 6.3 | **0.012** | 0.1 | 0.773 |
| Off. M x **Par. M** x **Par. W** | 1 | 6.0 | **0.014** | 0.2 | 0.679 | 0.2 | 0.650 | 0.3 | 0.617 | 0.0 | 1.000 | 2.4 | 0.124 | 2.9 | 0.088 | 8.7 | **<0.01** | 7.2 | **<0.01** | | | | |
| Off. W x **Par. M** x **Par. W** | 1 | 0.2 | 0.648 | 0.0 | 0.842 | 2.6 | 0.105 | 0.2 | 0.683 | 1.1 | 0.289 | 1.0 | 0.324 | 0.0 | 0.968 | 1.2 | 0.267 | 0.4 | 0.532 | 1.9 | 0.169 | 0.1 | 0.718 |
| Off. M x Off. W x **Par. M** | 1 | 2.1 | 0.145 | 3.9 | **0.048** | 0.6 | 0.426 | 0.2 | 0.632 | 3.4 | 0.063 | 1.7 | 0.186 | 2.5 | 0.115 | 5.6 | **0.018** | 4.3 | **0.038** | | | | |
| Off. M x Off. W x **Par. W** | 1 | 0.0 | 0.885 | 0.1 | 0.796 | 1.2 | 0.267 | 0.9 | 0.352 | 1.1 | 0.289 | 1.0 | 0.313 | 4.9 | **0.026** | 1.7 | 0.199 | 2.6 | 0.105 | | | | |
| Off. (M x W) x **Par. (M x W)** | 1 | 0.6 | 0.429 | 1.2 | 0.281 | 1.2 | 0.269 | 0.2 | 0.650 | 0.3 | 0.596 | 1.6 | 0.212 | 1.1 | 0.288 | 0.6 | 0.432 | 1.4 | 0.238 | | | | |

**Table S3: Summary of the linear mixed-effect model for main and interaction effects of offspring and parental treatments (the latest highlighted in bold) on adult offspring.** Degrees of freedom followed by χ2 values and P values are given for all the effects analysed. Significant results are shown in bold, and the colour indicates the direction of the effect (positive in blue, negative in red)

| | | Plant traits | | | | | | | | | | | | | | | | AMF colonization | | | |
|---|---|---|---|---|---|---|---|---|---|---|---|---|---|---|---|---|---|---|---|---|---|
| | | Whole-plant traits | | | | Leaf traits | | | | | | | | Root traits | | | | | | | |
| Source of variation | df | Total biomass | | Root mass factor (RMF) | | Leaf dry matter content (LDMC) | | Specific leaf area (SLA) | | Leaf P content | | Leaf C:N ratio | | Specific root length (SRL) | | Average root diameter | | % Fine roots | | Root length colonized by AMF (%RLC) | | Arbuscules: vesicles ratio | |
| | | χ2 | P | χ2 | P | χ2 | P | χ2 | P | χ2 | P | χ2 | P | χ2 | P | χ2 | P | χ2 | P | χ2 | P | χ2 | P |
| Seed mass | 1 | 0 | 0.99 | 0.1 | 0.8 | 0 | 0.85 | 1 | 0.32 | 1.1 | 0.29 | 2.7 | 0.10 | 3.3 | 0.07 | 2.6 | 0.10 | 0.1 | 0.80 | 5.5 | **0.029** | 0 | 0.986 |
| Offspring Mycorrhizal inoculation (Off. M) | 1 | 23.9 | **<0.01** | 7.9 | **<0.01** | 15.5 | **<0.01** | 0 | 0.918 | 471.5 | **<0.01** | 63.8 | **<0.01** | 581.8 | **<0.01** | 143.1 | **<0.01** | 301.7 | **<0.01** | | | | |
| Offspring Water availability (Off. W) | 1 | 127.6 | **<0.01** | 27.4 | **<0.01** | 69.6 | **<0.01** | 56.4 | **<0.01** | 1.9 | 0.17 | 34.7 | **<0.01** | 41.4 | **<0.01** | 8.1 | **<0.01** | 30.3 | **<0.01** | 0 | 0.938 | 13.1 | **<0.01** |
| **Parental Mycorrhizal inoculation (Par. M)** | 1 | 0.6 | 0.422 | 0.4 | 0.506 | 0.1 | 0.747 | 0.8 | 0.366 | 1.5 | 0.225 | 0.3 | 0.586 | 0 | 0.983 | 0.3 | 0.598 | 1.8 | 0.178 | 0 | 0.868 | 0.1 | 0.731 |
| **Parental Water availability (Par. W)** | 1 | 1.7 | 0.19 | 0.2 | 0.665 | 0.9 | 0.336 | 0.4 | 0.548 | 2 | 0.156 | 4.0 | 0.046 | 1 | 0.313 | 0.5 | 0.464 | 1.2 | 0.272 | 0.3 | 0.587 | 1.3 | 0.248 |
| Off. M x Off. W | 1 | 254.7 | **<0.01** | 44.6 | **<0.01** | 145 | **<0.01** | 122.8 | **<0.01** | 7.9 | **<0.01** | 67.7 | **<0.01** | 51.6 | **<0.01** | 0 | 0.895 | 0 | 0.897 | | | | |
| **Par. M x Par. W** | 1 | 0.4 | 0.506 | 0.8 | 0.378 | 0.2 | 0.651 | 0.1 | 0.76 | 0.1 | 0.709 | 1.3 | 0.252 | 1.3 | 0.262 | 2 | 0.16 | 0.6 | 0.443 | 0.6 | 0.447 | 0.3 | 0.588 |
| Off. M x **Par. M** | 1 | 0.6 | 0.439 | 0 | 0.845 | 0.6 | 0.441 | 0.6 | 0.453 | 0 | 0.84 | 0.4 | 0.510 | 0 | 0.827 | 1.5 | 0.228 | 1 | 0.307 | | | | |
| Off. M x **Par. W** | 1 | 0.2 | 0.69 | 0.5 | 0.495 | 0.1 | 0.707 | 0.1 | 0.722 | 2.3 | 0.127 | 0.1 | 0.750 | 0.8 | 0.358 | 0.6 | 0.445 | 0.2 | 0.641 | | | | |
| Off. W x **Par. M** | 1 | 1 | 0.322 | 1.3 | 0.26 | 5.0 | **0.026** | 2.5 | 0.111 | 0.4 | 0.509 | 3.1 | 0.078 | 0 | 0.891 | 0.1 | 0.803 | 0.3 | 0.57 | 5.8 | **0.016** | 0.1 | 0.789 |
| Off. W x **Par. W** | 1 | 1.3 | 0.256 | 0.1 | 0.795 | 0.2 | 0.642 | 0 | 0.942 | 0.3 | 0.591 | 0.8 | 0.378 | 0.1 | 0.732 | 0.9 | 0.331 | 0 | 0.966 | 0 | 0.933 | 0 | 0.981 |
| Off. M x **Par. M** x **Par. W** | 1 | 0.5 | 0.485 | 1.1 | 0.305 | 0.2 | 0.69 | 0.2 | 0.694 | 1.6 | 0.211 | 0.2 | 0.672 | 0.2 | 0.642 | 0.3 | 0.616 | 0.6 | 0.433 | | | | |
| Off. W x **Par. M** x **Par. W** | 1 | 0 | 0.879 | 0 | 0.882 | 3.7 | 0.056 | 2.3 | 0.128 | 1.5 | 0.224 | 0.1 | 0.750 | 1.6 | 0.204 | 0.2 | 0.674 | 0.4 | 0.536 | 0 | 0.861 | 2 | 0.153 |
| Off. M x Off. W x **Par. M** | 1 | 0 | 0.969 | 0.4 | 0.545 | 1.5 | 0.214 | 1.8 | 0.182 | 0.5 | 0.487 | 0.1 | 0.765 | 0 | 0.996 | 0.9 | 0.353 | 0 | 0.942 | | | | |
| Off. M x Off. W x **Par. W** | 1 | 0 | 0.924 | 0 | 0.893 | 0.7 | 0.404 | 0.2 | 0.632 | 2.1 | 0.151 | 0.0 | 0.894 | 0.4 | 0.504 | 0.3 | 0.602 | 0 | 0.969 | | | | |
| Off. (M x W) x **Par. (M x W)** | 1 | 0.2 | 0.675 | 1.1 | 0.301 | 0.2 | 0.629 | 0.1 | 0.717 | 2.5 | 0.113 | 0.0 | 0.901 | 1.3 | 0.253 | 0 | 0.849 | 0.1 | 0.776 | | | | |

20
